# Collective chemotaxis in a Voronoi model for confluent clusters

**DOI:** 10.1101/2021.11.15.468718

**Authors:** E. Lawson-Keister, M. L Manning

## Abstract

Collective chemotaxis, where single cells cannot climb a biochemical signaling gradient but clusters of cells can, has been observed in different biological contexts, including confluent tissues where there are no gaps or overlaps between cells. Although particle-based models have been developed that predict important features of collective chemotaxis, the mechanisms in those models depend on particle overlaps, and so it remains unclear if they can explain behavior in confluent systems. Here, we develop an open-source code that couples a 2D Voronoi simulation for confluent cell mechanics to a dynamic chemical signal that can diffuse, advect, and/or degrade, and use the code to study potential mechanisms for collective chemotaxis in cellular monolayers. We first study the impact of advection on collective chemotaxis, and delineate a regime where advective terms are important. Next, we investigate two possible chemotactic mechanisms, contact inhibition of locomotion and heterotypic interfacial tension, and demonstrate that both can drive collective chemotaxis in certain parameter regimes. We further demonstrate that the scaling behavior of cluster motion is well-captured by simple analytic theories.

**Statement of Significance:** The ability of cell collectives to respond to biochemical signals, called collective chemotaxis, is crucial for many important processes including embryonic development and wound healing. We developed an open-source computational model that couples biochemical signaling gradients to confluent cell layers, where there are no gaps between cells. Our model demonstrates that two experimentally observed local cell behaviors – neighbor-induced changes to interfacial tension or a tendency of cells to repel one another after they come into contact – can drive collective chemotaxis. We also highlight a regime in which the motion of migrating cells can alter the gradient.

## 2 Introduction

Many cell types exhibit chemotaxis, where they migrate up a chemical gradient. Chemotaxis is seen in a variety of different biological processes such as wound healing [1, 2, 3], cancer metastasis [4, 5, 6, 7], and development [8, 9, 10, 11, 12, 13]. Different cell types sense and respond to a chemical gradient in different ways; some cell types can sense the change in concentration across their length, which gives them explicit directional information about the chemical gradient [14, 15, 16, 17]. However, other cell types only sense the average signal strength at their location and must use other mechanisms to climb the gradient [18]. There are even some cells that can sense and migrate up steep gradients but require other mechanisms to respond to situations with shallower gradients [19, 20].

One strategy cells use when they cannot chemotax alone is to use interactions between neighboring cells to sense and then collectively migrate up the gradient as a cluster, in a process called collective chemotaxis [21, 22]. One example of this is the neural crest cells in the *Xenopus* embryo, in which clusters of cells will climb a *stromal cell-derived factor 1* gradient but single cells will not [9]. While the exact mechanism driving this behavior is still unknown, contact inhibition of locomotion [23], cluster confinement [24], and asymmetric actomyosin contraction [25] are implicated as possible mechanisms. Additionally, when lymphocytes are exposed to ligand gradients they will form clusters that can consistently climb the gradient. Interestingly, depending on the strength of the gradient, single cells will either not respond in the case of shallow gradients or migrate in the opposite direction of clusters in steep gradients [21]. Similarly, inside the Drosophila ovary, border cells migrate collectively up a ligand gradient by integrating the difference in signal levels between cells in a group [22, 26, 27, 28, 29, 30, 31].

While the mechanism for collective chemotaxis is not known in every case, contact inhibition of locomotion (CIL) is commonly implicated [32, 33, 23, 34]. Contact inhibition of locomotion is a well-documented and extremely common cellular behavior, not limited to chemotaxis [35, 36, 37, 38, 39], whereby cells stop moving toward one another once they come into contact. In some cases, CIL occurs because cells lack the cytoskeletal and adhesion machinery to migrate along the surface of another cell [40], while in other cases cell-cell contacts trigger a signaling cascade that inhibits migratory behavior [41].

A recent theoretical and computational paper [42] investigated a model where individual cells are modeled as repulsive particles and demonstrated that CIL can indeed induce collective chemotaxis up the gradient even when individual cells cannot sense the gradient. In this model, particles that overlap experience a change to their direction of migration that reduces the overlap. The authors showed that if the magnitude of this change was proportional to the average biochemical signal strength at that location, then the combination of all such interactions within the cluster would drive the cluster up the gradient. This is a simple and robust mechanism for collective chemotaxis that captures many features seen in experiments.

An important open question, however, is whether this mechanism is restricted to cell types that are well-modeled as overlapping particles. There are many cell types that do not remain spherical, and instead change their cell shape dramatically when interacting with other cells, such as epithelial cells in confluent tissues. Moreover, in bulk systems, the collective behavior of particle-based and confluent models are quite different. In particulate models, the fluid-to-solid transition is driven by an increase in particle overlaps with increasing density or packing fraction [43, 44, 45], which is known as jamming [46, 47]. In contrast, in confluent vertex and Voronoi models, the packing fraction is always unity and the fluid-solid transition is driven by a geometric incompatibility, such as changes to the shape of cells in the tissue [48, 49, 50, 51]. The vertex model also exhibits topological cusps in the energy landscape which contribute to interesting nonlinear responses in certain regimes [52, 53]. Given such differences, it is not obvious whether CIL can induce collective chemotaxis in a confluent model.

To address this question, we study simulations of a Voronoi model for 2D biological tissues coupled to a concentration gradient with independent dynamics, where individual cells can only sense the average concentration at their location (Fig. 1). We study several possible mechanisms that could drive collective chemotaxis in such systems, including a version of CIL that is well-defined in a confluent monolayer, as well as a mechanism based on interfacial tension between two different cell types. We find that both mechanisms are capable of driving collective chemotaxis and highlight specific experiments that could distinguish between the two mechanisms.

**Figure 1:**
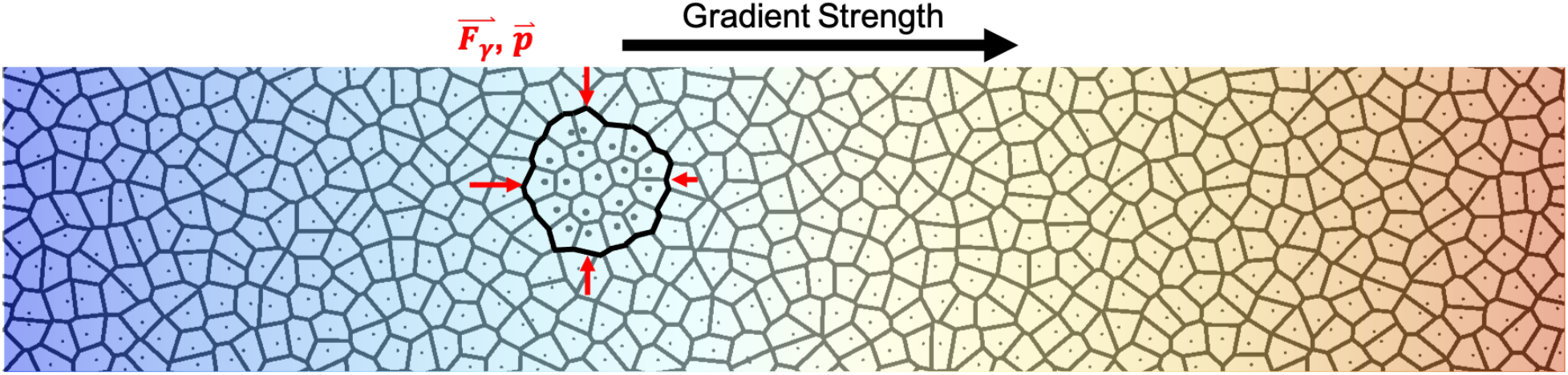
A confluent tissue is composed of exterior cells and cluster cells with an interfacial tension between the two types, denoted by a thick dark line. The cells in the cluster respond to chemical signaling and change either their i) CIL or ii) HIT with respect to the exterior cells. Exterior cells do not respond to the gradient. Due to a higher concentration of the signal on the front of the cluster, there is either a net polarity or force across the cluster driving it up the signaling gradient.

## 3 Methods

We simulate confluent monolayers using a Voronoi model which, along with vertex models, has been identified as a good model for confluent tissues [50, 54, 55, 56, 57, 58, 59]. Each cell, *i*, is defined by a Voronoi tesselation of the cell centers and is denoted by a polygon of a given area, *A_i_*, and perimeter, *P_i_*.

Then the energy functional for the *i*th cell is described by forces due to intercellular interaction:

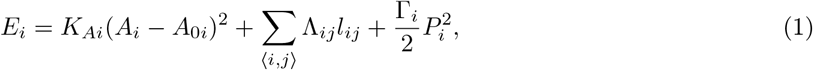

where the mechanical interaction forces between cells are given by *F_i_* = −∇_*i*_*E*. The first term in the energy describes the volume incompressibility of the cell. By assuming the lateral modulus of the cells is high so that height fluctuations along the monolayer are small and the actin-myosin ring is acting on the apical side, then one can show that the volume incompressibility is approximated at first order by a quadratic penalty on the area of the apical face *A_i_* when it deviates from the preferred area *A*_0*i*_, with modulus *K_Ai_*. The second term, Λ_*ij*_, represents the competition between adhesion due to molecules such as cadherins which try and elongate interacting edges, *l_ij_*, and line tension which tries to shrink these edges. The third term describes a non-linear perimeter restoring force that could be caused by an assortment of phenomena such as induced surface elasticity from the actomyosin ring [54] or a finite pool of adhesion molecules along the cell membrane which limits the perimeter.

Eq 1 can be further simplified by assuming that each cell has the same line tension, perimeter contractility, and area and perimeter moduli, *K_A_* and *K_P_*, respectively. Then by writing the energy functional in terms of cell perimeters *P_i_* and preferred perimeter *P*_0_ we find an energy functional that depends only on the cell shape.

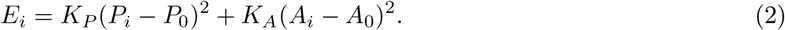

Additionally, we will be examining a mixture of two different cell types, cluster cells and external cells. As in previous work [52, 60], we assume these cells experience heterotypic interfacial tension (HIT) in which they recognize neighbors of different cell types and experience an additional energy cost for contacting the opposite type:

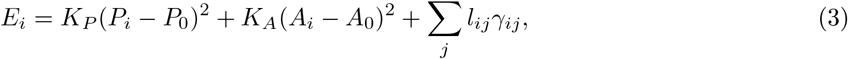

where *γ_ij_* represents the additional interfacial tension between the two cell types. This rule has been shown to create sharp but deformable interfaces [52]. Additionally, this will cause compartmentalization between the two cell types [60] which maintains the boundary between the cluster cells and external cells. The model can be nondimensionalized by expressing all lengths in units of 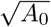 and all time in units of *τ_ch_* = 1/(*μK*_*a*_*A*_0_). The integration time step is set to *dt* = 0.001 and we set all cells to be fluid-like with *P*_0_ = 3.85 and *A*_0_ = 1. Consistent with previous work, we set *K_P_* = 1, *μ* = 1, and *K_A_* = 100 to ensure that cluster cells maintain their area even under large compression from HIT.

The final ingredient to the model is a biochemical signaling gradient. The system will be overlaid by a scalar field representing the biochemical gradient which evolves according to an advection-diffusion equation. In many biological contexts, enzymatic activity degrades signaling at a roughly constant rate, which contributes to an additional degradation term [61, 62]. Together, this leads to the following evolution equation for the scalar concentration 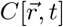 of a biochemical signal:

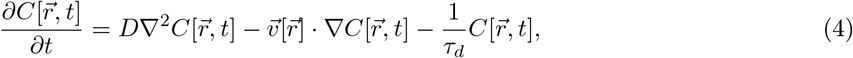

where *D* is the diffusion coefficient for the chemical, 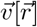 is the velocity of the cell at position 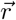, and *τ_d_* is the characteristic degradation time of the chemical. In this paper, we focus on a simple geometry with a line of source at the top of the system and a sink at the bottom and assume no degradation of the signal. This generates a linear gradient in steady state. However, it should be noted that the simulation tool we develop here can accurately evolve from any initial conditions and with any degradation parameter enabling the study of more complicated gradients.

We will assume a homogeneous, effective diffusion constant *D* that averages over smaller-scale features such as cell membrane permeability. As we are focused on the behavior of cells that can not individually climb a gradient, we further assume that cells are able to estimate the average signal across their area – in other words, the absolute concentration at their location – but that they cannot calculate other features of the signal, such as the local gradient. They then use this average signal to alter their individual properties. The code is available for download at https://github.com/Manning-Research-Group/ClusterVoronoiCode. Additional details of the simulation code are described in the supplementary material.

## 4 Results

### 4.1 The role of advection

Before coupling the cell mechanics to the signaling gradient, we investigate the importance of advection. As individual cells migrate through the tissue, it is possible that they may drag signaling molecules along with them and alter the local concentration. Whether this is a significant effect depends on the competition between cell-motion-driven advection and diffusion, described by the Peclet number:

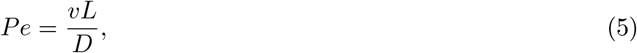

where *v* is the velocity of the advective flow, *L* is the characteristic length, and *D* is the diffusion coefficient. Thus for higher Peclet numbers (*Pe* ≫ 1) advection dominates, while for low Peclet numbers (*Pe* ≪ 1) diffusion dominates. Since the bare diffusion constant for most biochemical signaling molecules inside a cell is much greater than the velocity of cells in the tissue [63], it is expected that diffusion will dominate. Therefore, many models do not account for the role of advection. However, for example, if a given cell type has a cell membrane that is largely impermeable to a given signaling molecule, it is possible that the effective diffusion constant could be quite small and compete with the advection timescale. Therefore, we first characterize advection in this system so that we can specify precisely when it can be neglected.

We simulate a cluster of cells in a steady state linear concentration gradient. The cells in the cluster are all being pulled by a body force so that in the overdamped limit they move at a velocity, 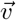, from a region of low concentration to a region of high concentration for different diffusion coefficients. Then we measure the total concentration *C_a_* inside the cluster when we include the advection term and compare it to the concentration *C_d_* in the purely diffusive case when we do not include that term:

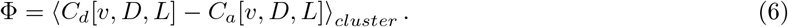

The top panel of Fig 2(A) is a schematic diagram of a moving cell cluster, and the bottom panel shows the difference in concentration Φ associated with that motion. Fig 2(B) shows the concentration of signal in the system as a function of *y*-position. The red(blue) line corresponds to the behavior when advective terms are neglected(included), highlighting that, as expected, the moving cluster of cells does entrain and drag along some of the chemical signal. The simulation data points in Fig 2(C) highlight that this average difference in concentration within the cluster increases with increasing Peclet number. It is worth noting that the velocity field constructed in simulations is discontinuous from one cell to another and, to avoid numeric instabilities, in simulations with high Peclet number we decrease our integration step significantly to *dt* = 10^−4^.

**Figure 2:**
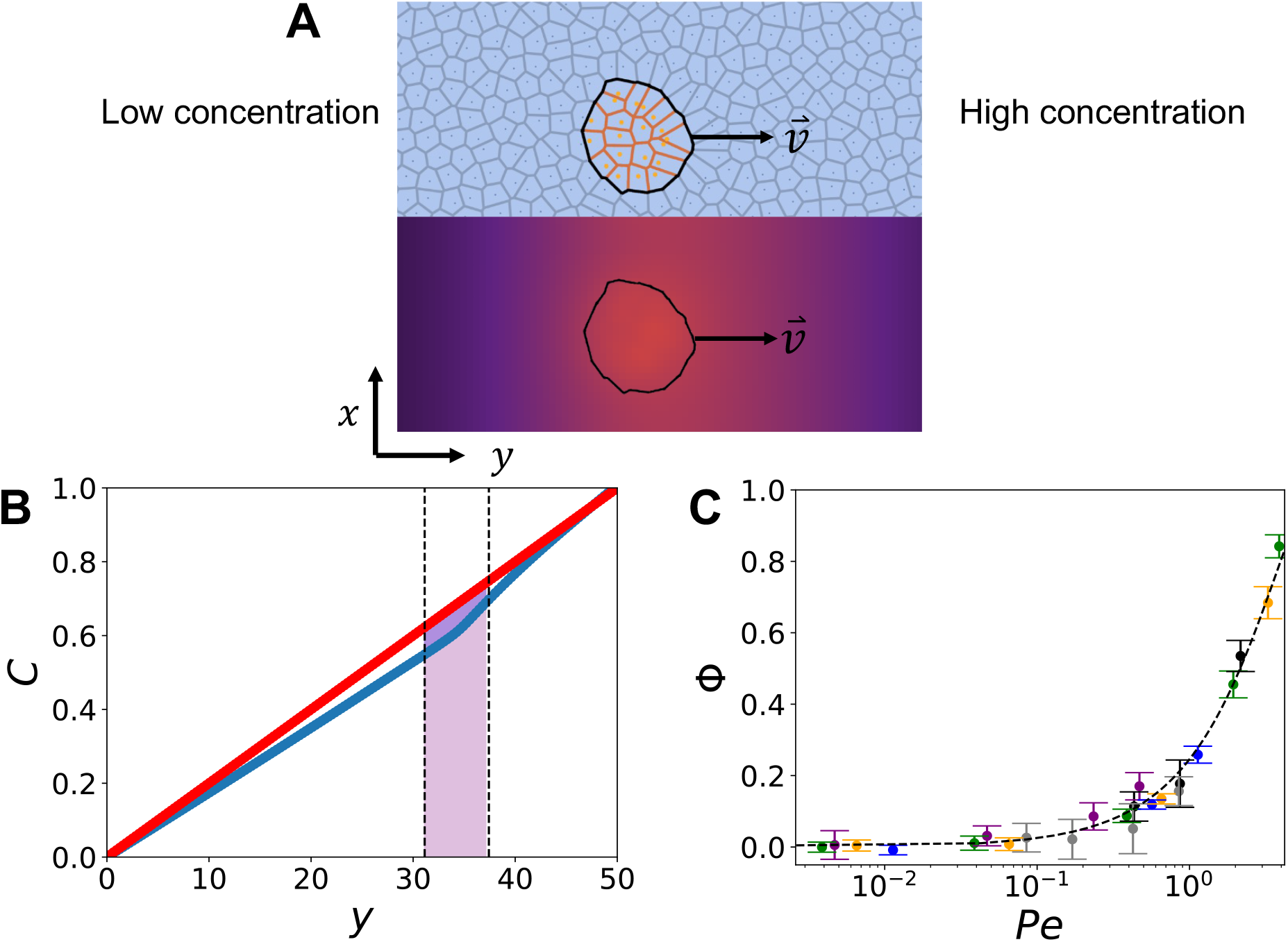
(A) Top panel: snapshot of a cluster moving up the gradient. Bottom panel: The change in the concentration gradient Φ due to advective terms. (B) A projection of the concentration in the system in the y-direction for *Pe* = 1. The dotted lines represent the location of the cluster. The blue line shows the concentration of signaling gradient in the presence of an advecting cluster and the red line shows the same gradient with pure diffusion. (C) The difference of the total concentration of signal inside the cluster with pure diffusion compared to the total concentration with advection Φ increases with increasing Peclet number (*Pe*). The cluster cells are climbing the gradient with various velocities: 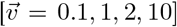 for the purple, green, orange, and blue points respectively over a range of diffusion constants between *D* = 10 and *D* = 10^4^. The grey and black points are for clusters with *N_c_* = 10, 30 respectively and 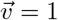. The dotted line is the predicted relationship from the toy model.

To predict the functional dependence of Φ on *Pe*, we develop a simple toy model. We assume a square cluster of side length *L* is climbing a linear gradient in the *y*-direction at a constant velocity 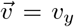, and therefore it will reach a steady state in which the rate that advection pulls the gradient with the cluster is equal to that lost by diffusion. Since the gradient is uniform in the *x*-direction, this generates a simple second-order differential equation:

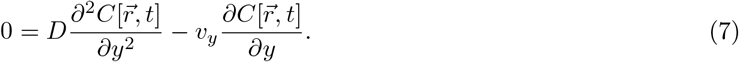

Integrating once we find that the gradient inside the cluster changes in *y* as a decaying exponential, where the integration constant is the change in gradient for the pure diffusion case:

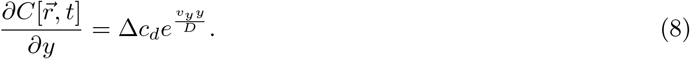

Integrating again and then applying the boundary conditions that the gradient is continuous across the cluster gives the concentration at every point inside the cluster:

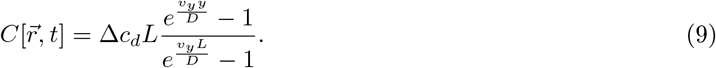

Then we can integrate a final time to find the total concentration inside the cluster in 1D. We multiply it by the length of the cluster to find the total signal inside a square cluster and subtract the total concentration inside the same cluster with just pure diffusion. This generates an analytical prediction for how the concentration scales with the Peclet number:

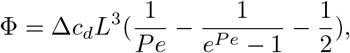

which is illustrated by the dashed line in Fig 2(C), showing that the prediction closely matches what we observe in our simulations. We show that this prediction holds over a range of cluster sizes, biochemical diffusivities, and cluster velocities. This demonstrates that the total concentration inside the cluster scales with the Peclet number itself rather than any of the independent parameters swept over individually. In both cases, we see that advection does not change the concentration of the cluster outside uncertainty until *Pe* ≈ 1, which is higher than what is observed in many experiments. For example, in the *Xenopus* embyro the cluster is around 100 *μm* with a velocity up the gradient around 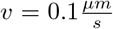 [25] within a sdf1 gradient which has a rough diffusivity of 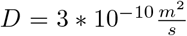[64] and yields a *Pe* ≈ 10^−2^, which is much smaller than what is predicted to cause noticeable advection. Therefore, for the remainder of the paper, we will neglect advection, though it may be interesting to revisit in the case of biochemical signals that diffuse very slowly.

### 4.2 Gradient-coupled contact inhibition of locomotion

Next, we will study collective chemotaxis in our confluent model. Previous work [42] demonstrated that collective chemotaxis emerges naturally when coupling a biochemical signaling gradient to a particle-based rule mimicking contact inhibition of locomotion (CIL).

Therefore, we first develop a rule for CIL similar to the one in Ref [42], that can be directly applied to a confluent model. Their rule was inspired by observations seen in various experimental systems including neural crest cells [23], rat kidney cells [33], and breast adenoacarcinoma cells [32]. As in previous self-propelled Voronoi, SPV, models [58, 59], we assume each cell *i* has polarity 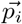 and experiences physical forces from the surrounding cells given by 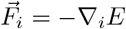. Then the cell’s motion will be over-damped:

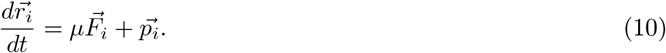

The direction of self-propulsion has its own dynamics:

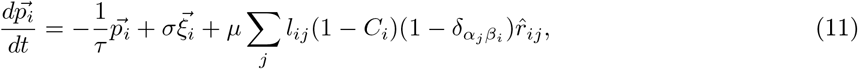

where *μ* is the inverse of the drag coefficient, *τ* is the self-propulsion persistence time for a single cell, and 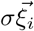 is Gaussian noise. The last term represents CIL, which alters the polarization so as to repel the two cells, and the strength of the repulsion depends on i) the concentration of signaling molecule and ii) how close the cells are to one another. Here *C_i_* = Δ*cy_i_* is the value of the concentration field at the center of cell *i*, which depends on its position *y_i_* and the change in gradient over a cell length Δ*c*. Then the direction of the change in polarity is 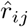, is the unit vector pointing from cell *j* to cell *i*. Whereas in previous work “cell closeness” was quantified by particle overlap [42], in our case cells are closer when they share a larger interface *l_ij_*. We choose *σ* = 1 such that an isolated cell has a velocity of unity. While difficult to actually quantify numerically the range of the polarization, we take inspiration from [9] which shows that the sdf1 gradient in neural crest cells causes asymmetry in Rac1 signaling, and therefore cell protrusion stability, across the cluster. This manifests in a Fluorescence Resonance Energy Transfer (FRET) efficiency ratio from the back to the front of the cluster between 0.5 – 0.9 [9]. Therefore, we choose our polarization such that the ratio between the CIL on the front the cluster to the back is in the same range.

In addition, since in our model we have cells surrounding the migratory cluster (while in previous work the surrounding environment was not directly modeled), we also have to specify cell types *α_j_* and *β_i_*. Interestingly, if the cells of the cluster experience the same magnitude of contact inhibition of locomotion with both the exterior cells and other cluster cells there is no chemotactic response, as we show in Supplementary Fig 9. Since many different cell lines experience contact inhibition of locomotion with specific types of cells [38], for the remainder of the paper we will assume that the cluster cells only experience heterotypic contact inhibition of locomotion between themselves and exterior cells. This rule is represented by the delta function in the last term of Eq 11.

In SPV models, a tissue composed of only one cell type, each cell’s velocity will be due to active forces from surrounding cells and the cell’s polarity. The cell’s polarity will relax to zero with persistence time *τ* but be driven away from zero by the noise. As illustrated in Fig. 3(A,B), when cells of different types share an edge the third term in Eq. 11 turns on. The cluster cell will experience a repulsive polarity away from the exterior cells (along the direction given by *dp*/*dt* in panel (A), which drives the polarization from *p_i_* towards *p_f_*) with a magnitude that depends on the concentration inside the cell and the length of the shared edge. Thus cells on the low-concentration side of the cluster, illustrated by the large red arrows in Fig 3(B), will experience a greater magnitude polarity inwards than the cells on the top of the cluster, illustrated by the small tan arrows in Fig 3(B).

**Figure 3:**
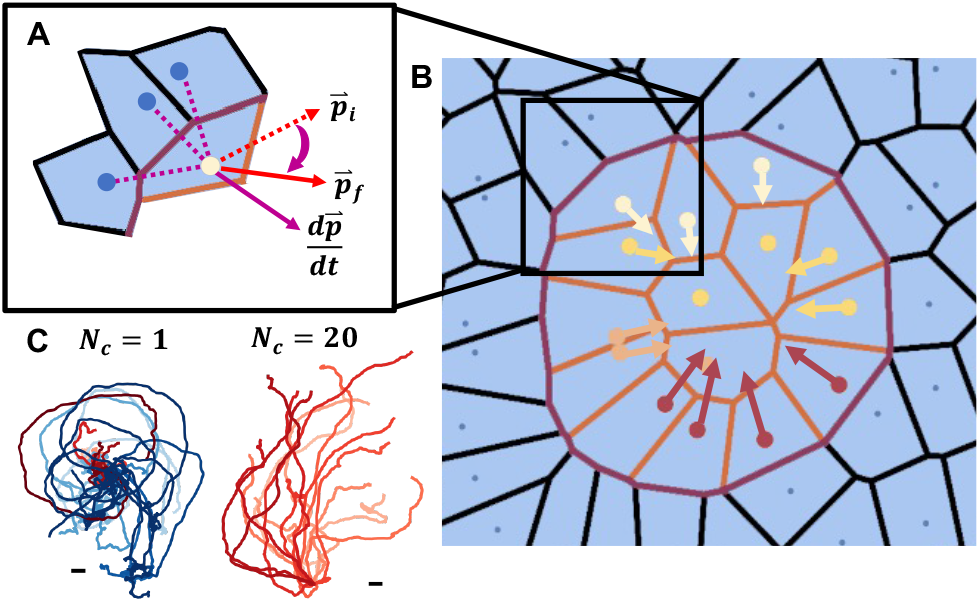
A.) The cluster cells experience heterotypic CIL as a change to their polarity away from external cells. The magnitude of this polarity corresponds to the magnitude of the gradient at that cell. B.) The entire cluster experiencing gradient-dependent CIL which generates net movement up the gradient. Lighter color cell centers represent cells that sense a higher gradient and therefore experience less CIL. C.) 20 Trajectories of Nc=1 and Nc=20 clusters. The scale bar represents one cell length. Red trajectories have final points on the higher-concentration side of the origin, while blue trajectories terminate in the lower-concentration half-plane.

Figure 3 (C) shows sample trajectories for single cells and clusters of cells with these CIL dynamics, demonstrating that single cells do not climb the gradient, but clusters do.

To quantify this effect more precisely, we measure the average velocity of the clusters up the gradient:

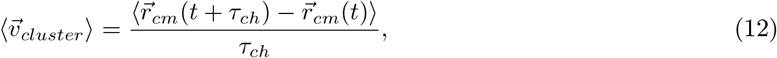

where 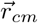 is the center of mass of the cluster and *τ_ch_* is the natural time unit. The inset to Fig. 4 shows that the velocity increases as cluster size increases, with a plateau after the cluster reaches around 10 cells. We see that this behavior holds over many persistence times *τ* and gradient slopes Δ*c*.

**Figure 4:**
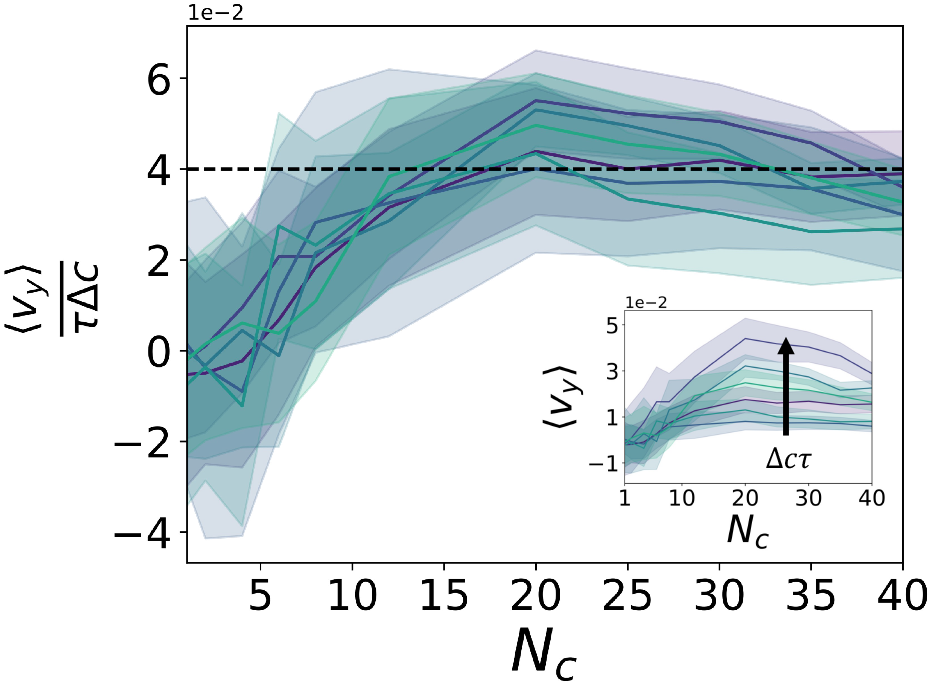
Inset: The average velocity of a cluster up the gradient. The velocity increases as cluster size increases and then plateaus after *N_c_* = 10. Main panel: The cluster velocity up the gradient collapses when correctly scaled by the persistence time and magnitude of the CIL. The dashed line represents predicted velocity from our toy model, which agrees after the cluster reaches *N_c_* = 8. In both panels, error bars are the average of the standard deviation of the cluster velocity in an individual trial.

To collapse the data, we attempt to predict the cluster velocity using a simple model. If we average the polarity of all the cells in the cluster over long times, we expect the contribution from the noise to average to zero. Assuming the polarity over the entire cluster will eventually reach a steady state we have:

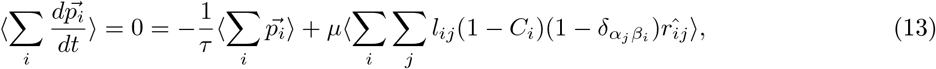

where we sum over each cell, *i*, in the cluster. We assume that the average polarity of the entire cluster is the average of the polarity of every cell in the cluster such that, 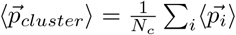. Then Eq. 13 can be simplified:

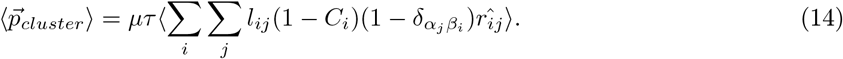

Then we will assume the cluster is roughly circular and turn this sum into an integral:

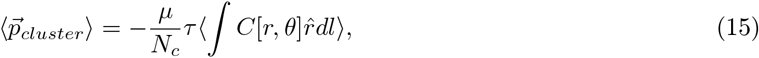

where *C*[*r*, *θ*] is the concentration inside the circular cluster and 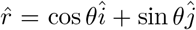 points radially outward from the center of the circle: *C*[*r*, *θ*] = *C_center_* − *r*Δ*c* sin*θ*, where *C_center_* is the concentration in the center of the cluster, and Δ*c* is the change in concentration over one cell length. Since each each cell has *A_i_* = 1 the radius of the cluster is 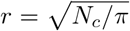. Integrating gives an expression for the steady state polarity of the cluster:

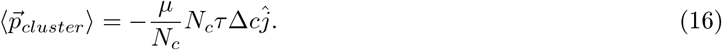

We see that we can collapse the velocity with the gradient slope Δ*c* and persistence time *τ* to a single curve which is independent of the number of cells in the cluster (Fig. 4). This matches closely with what was seen in lymphocytes exposed to ligand gradients in which above a certain cluster size, around *N_c_* = 20 in their experimental system, the velocity is largely independent of cluster size [21]. This trend was also seen in *Xenopus* neural crest cells in which the cluster speed reported was independent of size [9]. Together this suggests that this mechanism is a reasonable model for collective chemotaxis in real systems.

Notably, the fluctuations of cluster velocity for small clusters are much larger than those for larger clusters. Since collective migration in biology can sometimes involve small clusters of cells and fluctuations might be biologically relevant, we next explore the source of those fluctuations in some detail. Visual observations of simulation dynamics highlight that the smaller clusters rotate or “turn” fairly frequently, presumably due to unbalanced torques generated during CIL, shown in the schematic diagram in Fig 5 (A). Moreover, once the clusters turn they move persistently until the polarity invoked by the CIL causes them to turn back to moving up the gradient. Large clusters, however, would seldom experience this turning.

**Figure 5:**
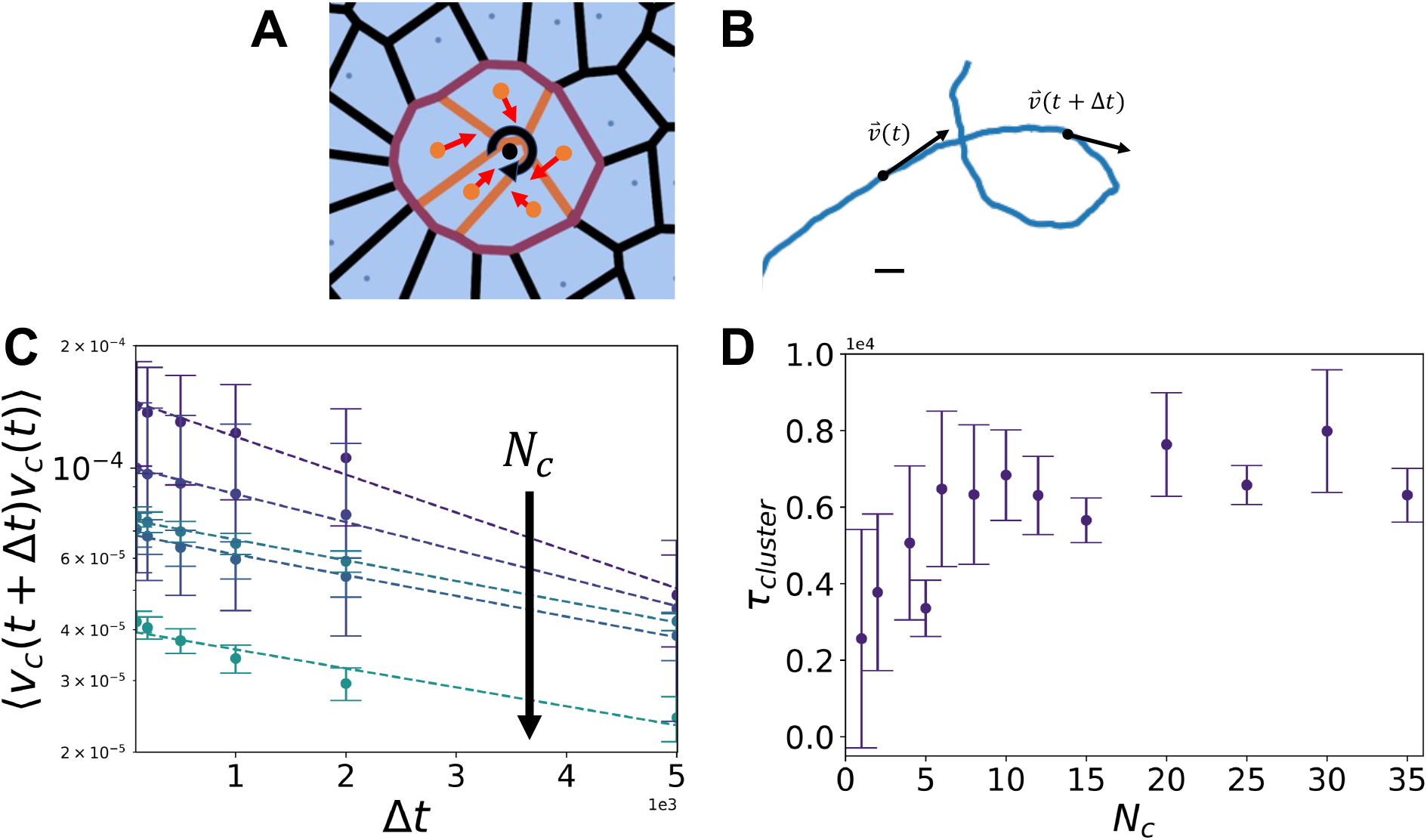
(A) The change in polarity from contact inhibition of locomotion, red, can cause a net torque, black, on small clusters. (B) The trajectory of a *N_c_* = 5 cluster as it turns. (C) The velocity autocorrelation function of various cluster sizes, *N_c_* = [4, 5, 10, 15, 25]. The dotted lines are the fits to exponential decay. (D) The persistence time of the cluster for various cluster sizes extracted from C.).

To validate these observations, we calculate the persistence time of clusters via a velocity autocorrelation function, as illustrated in the schematic diagram in Fig 5(B). For persistent random particles, we expect that the velocity autocorrelation should follow:

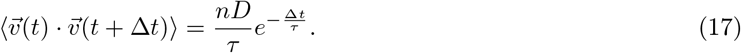

Indeed, a plot of the autocorrelation vs. Δ*t*, shown in Fig 5(C), is well-fit by an exponential decay and we extract a persistence time. Fig. 5(D) shows the statistics of persistence times as a function of cluster size, indicating that the persistence time of the cluster increases as the cluster size gets larger until around *N_c_* = 5 where it plateaus. This is reminiscent of a recent work looking at the persistence time of Vicsek aligning Brownian particles [65]. In that paper, the persistence time of clusters increases linearly with the size of the cluster. For a larger cluster to change direction, more individual cells are required to change direction. However, when a new cell which aligns at a much slower rate is added, the entire cluster’s persistence time decreases. We see for small cluster sizes the persistence time of the cluster increases linearly with cluster size in agreement with [65], but this effect eventually saturates. In 2D, *N* ~ 5 is when each additional new cell added to the cluster has a strong possibility of being added to the interior, without any exterior interface, and thus does not experience any CIL. These cells act like the less persistent cells in [65] as they primarily just add noise to the cluster’s polarity. There is a balance as the clusters become larger; they become more resistant to fluctuations from each newly added cell, experience a larger change in gradient over the cluster as the radius expands, and more noise is added by new internal cluster cells without significantly increasing the interface with cells on the exterior. It should be noted that we are only investigating the limit of high interfacial tension between the cluster cells and the exterior cells. We found that as the exterior surface tension becomes small, the clusters tend to completely break apart and thus have no preferred cluster size other than unity. So, while this balance in cluster size plays an important role in persistence time, the cluster does not seem to be able to self-select an optimal size using this mechanism.

### 4.3 Gradient-coupled HIT

The simulation tools we have developed also allow us to investigate other candidate mechanisms, in addition to CIL, for collective chemotaxis. Along with contact inhibition of locomotion, asymmetric actomyosin contractility has been implicated in the collective chemotaxis of neural crest cells [25]. Additionally, Eph-Ephrin ligand signaling has been associated with both collective migration of neural crest cells [66] and heterotypic interfacial tension [67]. In the Drosophila ovary, differences in mechanical tension between the front and back of the cluster drive polarization in the chemotactic border cells [68]. In a 3D Vertex model, polarized interfacial tension is sufficient to cause collective cluster migration [69]. Taken together, this suggests an alternative plausible mechanism for gradient climbing may be directly coupling HIT to a biochemical signaling gradient, and we develop a set of simulations to test this hypothesis.

Since this mechanism does not depend on self-propulsion, for simplicity, we assume cells follow over-damped Brownian motion such that 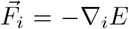 and where active fluctuations are Gaussian and governed by an effective temperature *T*:

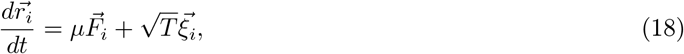

where the energy function is the same as described in Eq. 3, except that additional interfacial tension, *γ_ij_* is coupled to the gradient:

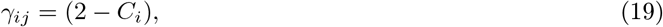

*C_i_* = Δ*cy_i_* where *C_i_* = Δ*cy_i_* is the value of the concentration field at the center of cell *i* which is based off the position of cell *i*, *y_i_*, and the change in gradient over a cell length Δ*c*. A schematic diagram of this type of interaction is shown in Fig. 6 (A,B). There must always be HIT between the cluster and exterior cells, even at the top of the gradient when *C_i_* = 1. While it is difficult to exactly measure heterotypic interfacial tension, there has been some work comparing the ratio in surface tensions between different cell types using cell doublet experiments in Zebrafish where the ratio between tensions is usually between 0.5 – 1 [70, 71].

**Figure 6:**
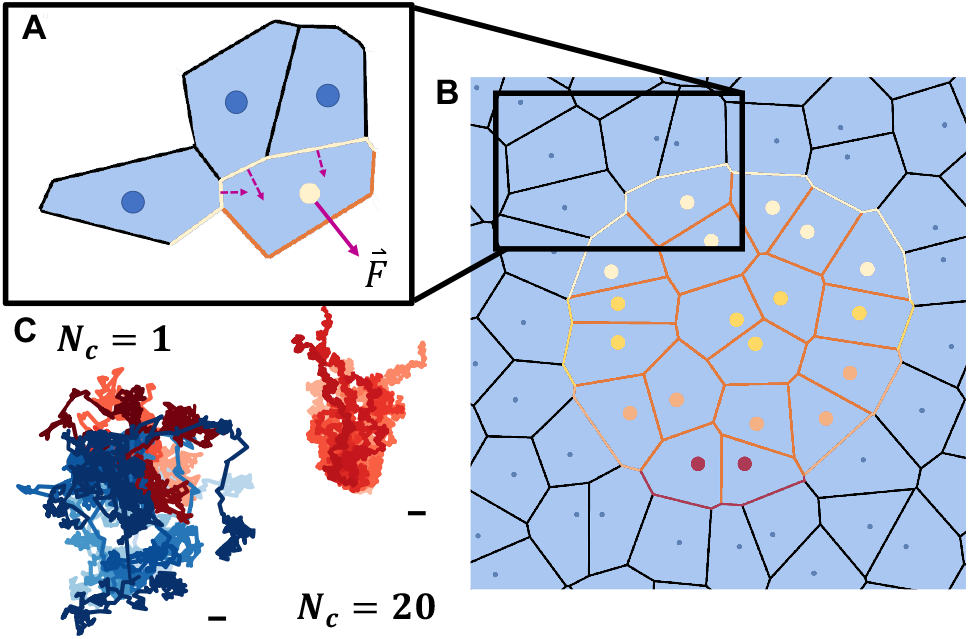
A.) The cluster cells experience heterotypic interfacial tension as a force away from the edges shared with exterior cells. The magnitude of this force corresponds to the magnitude of the gradient at that cell. B.) The entire cluster experiencing gradient-dependent HIT in which there is higher HIT on the rear of the cluster which propels the cluster up the gradient. Lighter color cell centers represent cells that sense a higher gradient and therefore experience less HIT. C.) 20 Trajectories of *N_c_* = 1 and *N_c_* = 20 clusters. The scale bar represents one cell length.

While it may be interesting to study even larger differences, we primarily look at values of Δ*c* such that the ratio of HIT on the front to the back of the cluster is between 0.7 – 1.

We expect that the force from the HIT should scale linearly with *γ_ij_* and point along the outward-oriented chord between the two touching cells 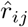 such that:

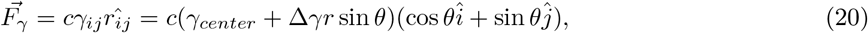

where *c* is a proportionality constant, as illustrated in Fig. 6(A).We assume this scaling, as we expect that the additional energy cost for heterotypic interfaces will generate a force on each cell pointing away from the interface and towards the center of the cluster. To validate this assumption we look at the average force on exterior cells in the cluster as a function of their position relative to the center in Supplementary Fig. 10; it matches this assumption. Since the net inward force is smaller on the high concentration side compared to the low concentration side of the cluster, we expect a net force on the cluster that drives the cluster towards regions of high concentration.

We simulate this model, resulting in sample trajectories as shown in Fig. 6(C). Similar to the CIL-coupled system, we see once again that clusters of cells collectively climb the gradient while individual cells do not. One obvious difference is that these trajectories are less persistent than for CIL; this mechanism is not as efficient at driving gradient climbing.

In addition, as shown in the inset to Fig. 7, by comparing clusters of different sizes we see that small clusters have a higher climbing velocity than larger clusters. Once again this can be explained using a simple model.

**Figure 7:**
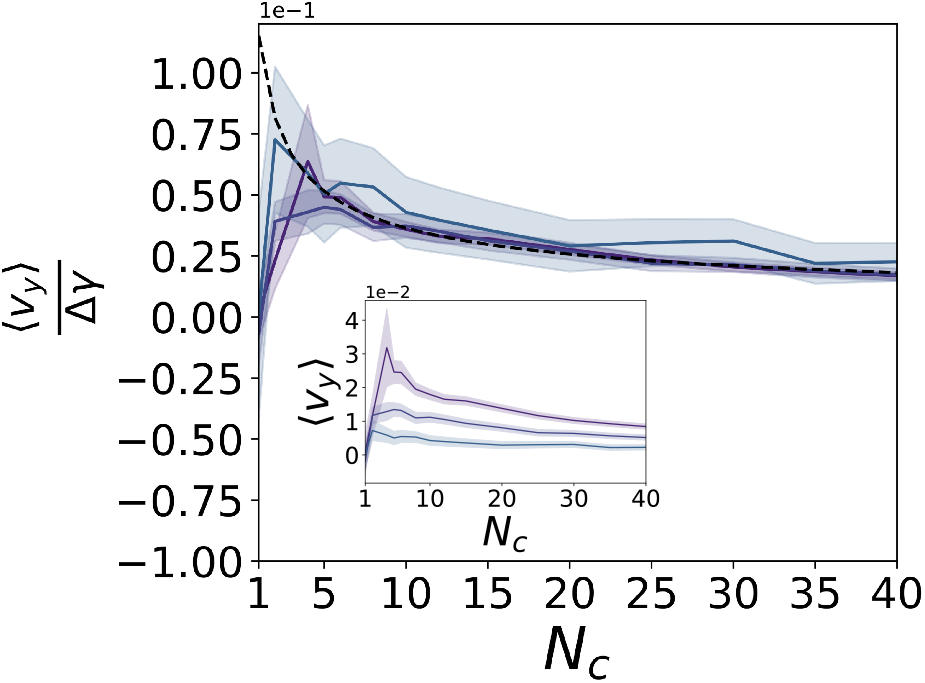
Inset: The average velocity of a cluster up the gradient. Main panel: The velocity collapses with the magnitude of the gradient’s effect on HIT (Δ*γ*) and follows the 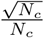 scaling predicted from the toy model shown by the dashed line.

We can perform the same integration steps as we did for the CIL-coupling Eq 14–16 to show that:

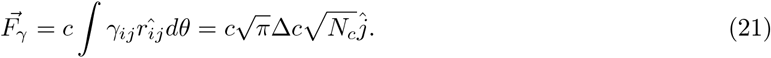

As shown in Supplementary Fig. 11, we find that the line tension force on the cluster observed in simulations follows this predicted functional form, and the best fit for the proportionality constant is *c* = 0.65. Just as before, if we average over many systems, we expect that the contribution for the Brownian noise and the non-gradient coupled forces to go to zero. Assuming that the force acting on the cluster is on average evenly distributed over the entire cluster, we have:

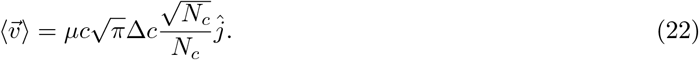

Using this functional form and the proportionality constant we identified previously, we can exactly predict the collapse of the average cluster velocity, shown by the dashed line in the main panel in Fig 7. Perhaps unexpectedly, as the cluster increases in size, the net velocity up the gradient decreases. This is due to the net force on the cluster being distributed over every cell in the cluster while only the outer cluster cells contribute to the gradient sensing. Therefore, we expect that as more cluster cells occupy the interior, the velocity up the gradient will decrease.

Notably, this scaling depends on *N_c_* while the gradient-coupled CIL does not. This occurs because in the gradient-coupled CIL *cell motion* is proportional to the length of heterotypic cell contacts as described in Eq. 11, while in the gradient-coupled HIT *the energy cost* is proportional to the length of heterotypic cell contacts. Thus, the force experienced is independent of this length.

This non-monotonic scaling with cluster size has been observed in other works as well. On one hand, an identical scaling of 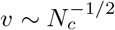 appears in a follow-up paper to [42] with an expansion on their 2D model, in which they investigate the role of both amplification and signaling within the cluster [72]. Strikingly, even though the mechanisms between our models are very different, they both lead to the same collective response with a mean force driving the cluster that scales like *N*^1/2^. This is driven primarily by polarity induced on the boundary of the cluster. On the other hand, in a different paper examining border cell migration in the Drosophila ovary, the authors predict that velocity would scale monotonically with cluster size in 2D but non-monotonically increase in 3D [27]. In that paper, the discrepancy in behavior is derived mainly from differences between 2D and 3D motion, how these differences manifest in terms of the forces that drive polarity on the cluster, and the dominant contribution to friction. Thus, delicate care must be put into each theory to replicate the behavior in the respective experimental system, as small differences in chemotactic response can lead to vastly different behaviors as a collective.

## 5 Discussion

We have created an adaptation of a Voronoi model [58, 59] for confluent tissues that allows individual cells to change their mechanical properties in response to the average biochemical signaling concentration at their location, and we have identified several local rules suggested by previous experimental work, that are able to drive collective chemotaxis.

We demonstrate that gradient-dependent contact inhibition of locomotion gives rise to collective chemotaxis, with smaller clusters climbing the gradient less efficiently than large clusters, up to a plateau. This, along with the previous results of CIL in particulate-based systems [42], suggests that CIL is a mechanism for collective chemotaxis regardless of the confluency of the tissue.

We also demonstrate a second possible mechanism for collective chemotaxis through a mechanism where the interfacial tension at the edge of the cluster depends on the local concentration, and in this case smaller clusters climb the gradient more efficiently than larger clusters.

This opens up an interesting question about how mechanisms of gradient climbing might influence the optimal size of collectively migrating clusters. In nature, clusters are often on the order of 20 cells, which suggests there may be a balancing between velocity and other considerations such as sensing error. Besides being able to gain more information about a gradient, cells will also use clusters to help reduce noise from signal processing [73]. In the future, it would be interesting to study how the mechanisms we present here in combination with cell sensing noise might generate an optimal cluster size.

Although we have focused here on two specific rules, the general, open-source code we make available with this manuscript will allow users to input any boundary conditions and chemical degradation rates to create an assortment of different steady state gradients.

Additionally, our code generally allows for cells to advect a biochemical signal. While we have focused here on situations where advection can be neglected (*Pe* < 1), we develop an analytic expression for how much advection changes concentration gradients in collective chemotaxis geometries. Since several experiments have shown that the effective diffusivity of some particles can be reduced by several orders of magnitude as they bind to the extra-cellular matrix [74, 75, 76] which could drastically increase the Peclet number in certain systems. This would make advection not only relevant, but crucial to modeling these non-steady state systems.

There may be other examples of processes where biochemical signaling can affect cell dynamics where this type of simulation and analysis could be helpful. For example, some tissues exhibit gradients in tissue fluidity [44], and it would be interesting to model the dynamic effect of coupling the cell shape and tissue fluidity to a signaling gradient. Another common mechanism for chemotaxis in cells is through run and tumble behavior [9, 77]. By coupling the biochemical gradient to the rotational diffusion of cells using a different equation for the polarity dynamics, we can investigate alternate climbing mechanisms.

Finally, the rheology of the tissue has been implicated in the ability of clusters to sense and react to gradients. There are conflicting theories that predict either that solid-like clusters [42] or fluid-like clusters are better sensors [73]. Clearly, the conclusions may depend on the precise model one uses for the mechanical interactions between cells. Therefore, one obvious extension to this work is to use models with the appropriate mechanical interactions for confluent tissues to investigate how different tissue fluidities affect the ability of cells to climb signal gradients.

## 6 Author Contributions

All authors contributed equally to this work.

## 7 Declaration of Interests

The authors declare no competing interests.

## 8 Acknowledgements

The authors acknowledge financial support from the Simons Foundation (#446222).

## 9 Supplemental Information

### 9.1 Computational modeling of the gradient

The signalling gradient is created by superimposing a scalar field on top of the prexisting cellGPU code [78]. The system is divided into a grid such that there are roughly 10 grid spaces inside each cell, 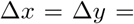 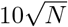, shown in Fig. 8. Initial conditions are set manually and currently are restricted to a source at the top of the gradient and a sink at the bottom.

**Figure 8:**
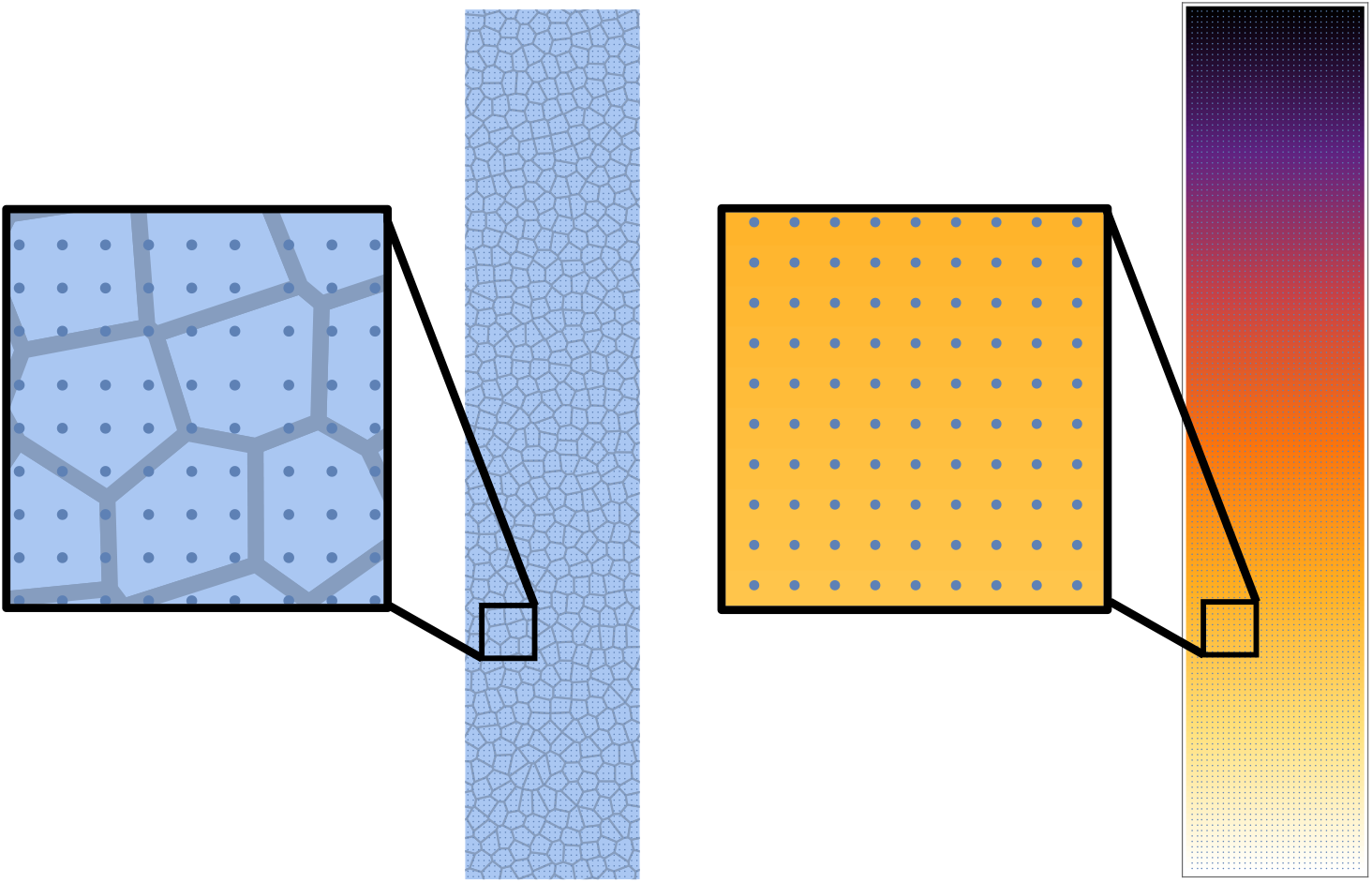
The scalar field representing the biochemical signal is divided into a grid that spans the system. The center of these grid points represents the location concentration in the grid square. Cells will average the concentration within their cell walls to determine the average signal at their location.

Then the gradient will evolve according to the advection-diffusion equation:

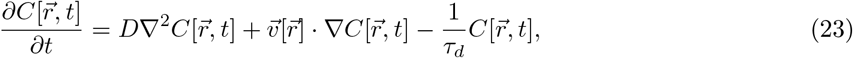

where 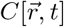 is a scalar field representing the concentration of the biochemical signal, D is the diffusion coefficient for the chemical, 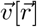 is the velocity of the cell at position 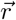, and *τ_d_* is the characteristic degradation time of the chemical.

This evolution is done using a simple central finite difference method such that:

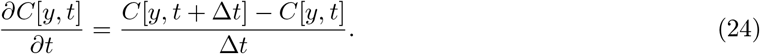

And similarly for second-order:

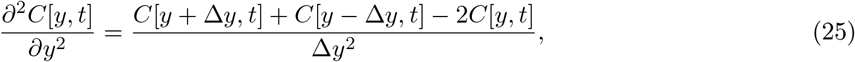

where Δ*t* = *dt* is our integration timestep and Δ*y* is the grid spacing.

At each time step, the cells will calculate a signal strength by taking the average concentration of each gridpoint within their cell walls. This is done by using a winding number algorithm such that if a gridpoint lies within a cell length of the center of a cell we check if that gridpoint lies within the cell walls. This is done by calculating the winding number Θ around the gridpoint which is the sum of the angles between two vertices of the cell and the gridpoint over all vertices on the cell.

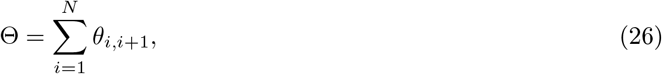

where *θ*_*i*,*i*+1_ is the angle between a vertex *i*, the next vertex in counterclockwise order *i*+1, and the gridpoint and *N* is the number of vertices of the cell. Such that if the winding number is a non-zero multiple of 2*π* the gridpoint lies within the cell or if the winding number is zero the gridpoint lies outside the cell.

This signal strength is then coupled to either the strength of CIL or HIT.

### 9.2 Homotypic CIL

In the main paper, we examine heterotypic CIL between cluster cells and exterior cells only but the cluster cells could also experience CIL between themselves. If we look at the case in which cluster cells experience CIL between other cluster cells and themselves and external cells and themselves we find that there is no net movement up the gradient (Fig. 9). Since the magnitude of the repulsive polarity is based purely on the average concentration of the cell, each cell will experience an isotropic repulsive polarity and thus will have a net-zero polarity due to CIL on average. However, altering the ratio between heterotypic (external cells with cluster cells) and homotypic (cluster cells with cluster cells) CIL, *κ*, we see that any difference between the two will recover the collective climbing behavior (Fig. 9).

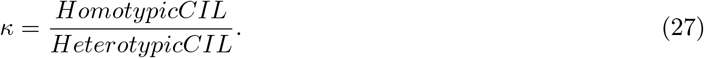

**Figure 9:**
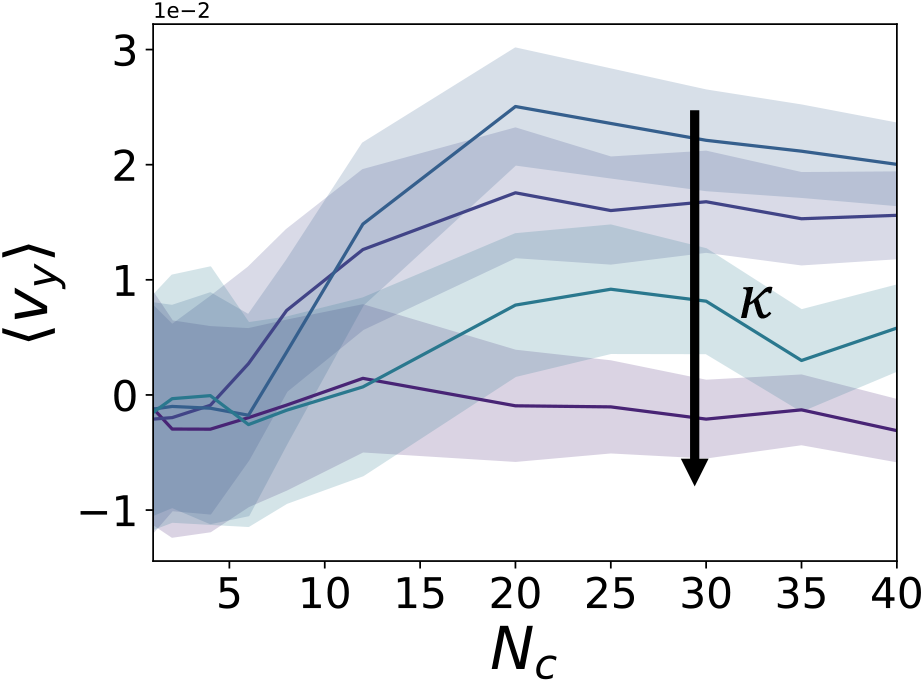
The cluster will not climb the gradient regardless of system size when CIL between all cells is the same magnitude (Purple line, *κ* = 1). But with any amount of difference in magnitude of homotypic and heterotypic CIL the cluster will experience collective chemotaxis

**Figure 10:**
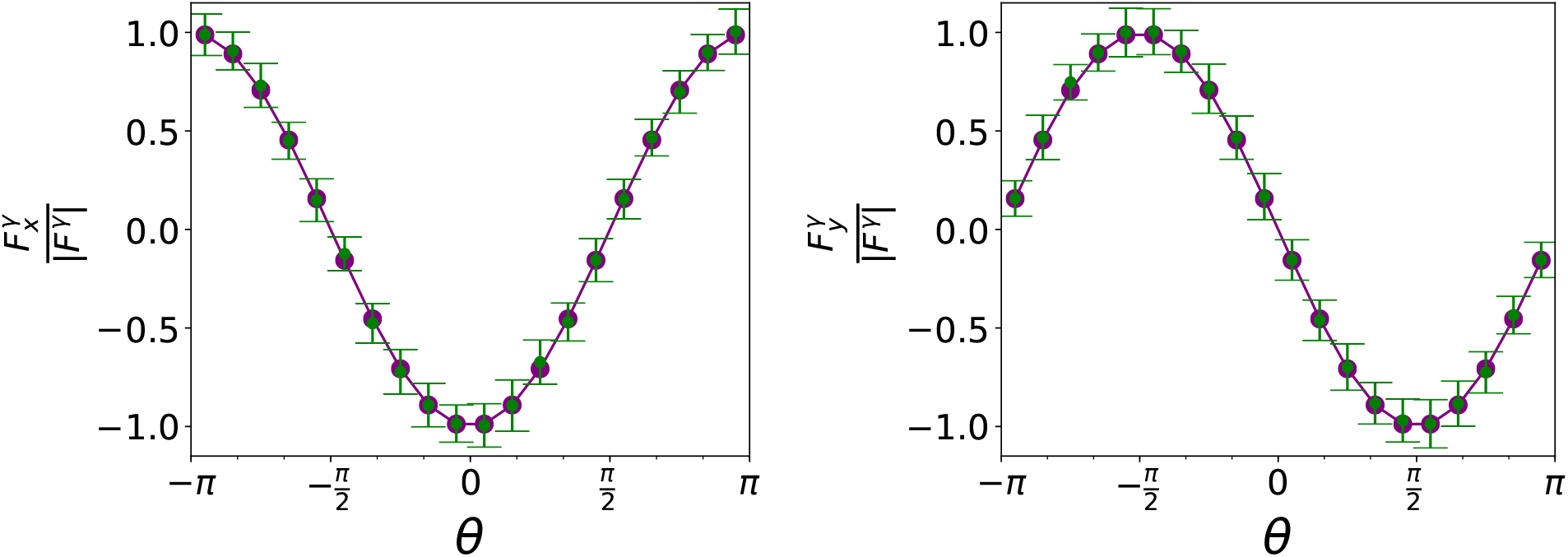
The normalized force from HIT on cells in the cluster as a function of cell orientation with respect to the center of the cluster. The green points are an ensemble average of the tension cluster cells experience while the purple points are what we assume in our theory for Eqn. 20

**Figure 11:**
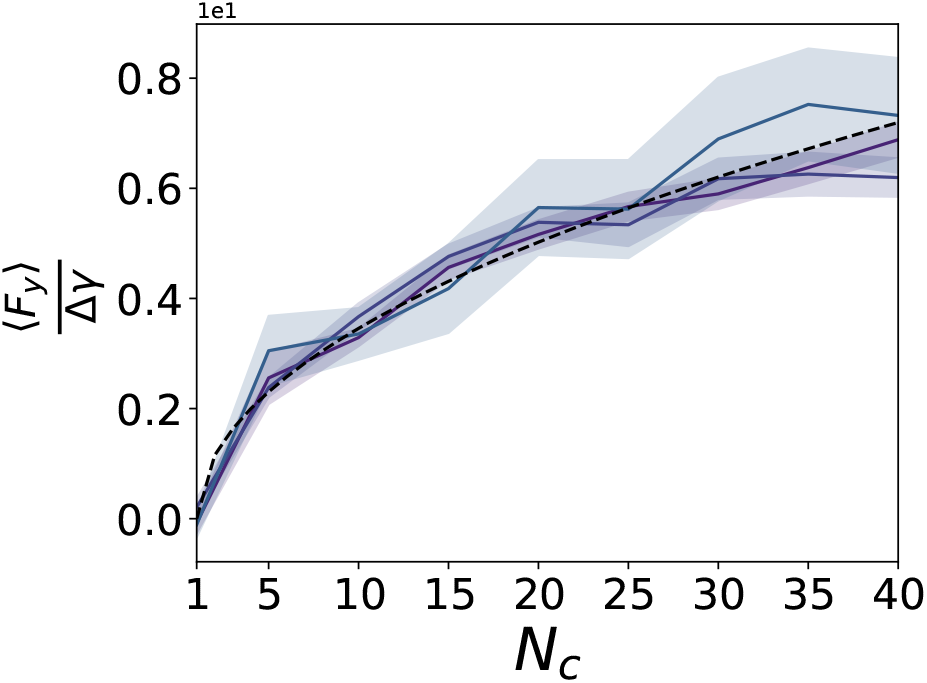
The average force on a cluster up the gradient due to the HIT. The force will collapse with the magnitude of the gradient’s effect on HIT Δ*γ* and follows the 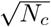 scaling from the toy model. We fit the proportionality constant *c* = 0.65

The magnitude of the climbing velocity will increase as *κ* increases as the magnitude of the polarity counteracting the heterotypic decreases. This means that pure CIL between just cluster cells and no interactions with external cells should also cause a collective chemotactic response but in the opposite direction.

## Notes

### Competing Interest Statement

The authors have declared no competing interest.

### Summary of Updates

We have added additional biological justification for the assumptions we made for both mechanisms. We have also provided an extended discussion of how our model results compare to other results in the literature including experimental data. We also added additional advection data in Figure 2.

